# Neuronal precursor cell persistence in Ganglioglioma is associated with ECM remodeling and immune cell infiltration

**DOI:** 10.64898/2026.04.18.719347

**Authors:** Jan Kueckelhaus, Lucas Hoffmann, Joëlle A Menstell, David Niklas Zimmer, Jasim Kada-Benotmane, Junyi Zhang, Juergen Beck, Oliver Schnell, Roman Sankowski, Philipp Sievers, Felix Sahm, Daniel Delev, Dieter Henrik Heiland

## Abstract

**Background:** Gangliogliomas (GGs) are low-grade glioneuronal tumors that frequently present with drug-resistant epilepsy. Although their indolent course contrasts with their high epileptogenic potential, the oncogenic mechanisms sustaining neuronal precursor-like populations within the tumor microenvironment remain poorly defined.

**Methods:** We performed spatial transcriptomic profiling on eight histologically confirmed GGs and matched healthy cortex to map the cellular and molecular architecture of the tumor microenvironment. Integrated analysis with weighted gene correlation network analysis (WGCNA) defined recurrent oncogenic programs and spatially resolved tumor–stroma interactions.

**Results:** Eight conserved gene modules emerged, encompassing physiological cortical, reactive glial, and oncopathological programs. The latter captured extracellular matrix (ECM) remodeling, vascular–immune signaling, and persistence of immature, proliferative neuronal-like states. Spatial modeling revealed that these oncopathological programs form structured niches at the tumor–brain interface, where radial glia–derived neuronal-like tumor cells coexist with immune and stromal elements engaged in ECM turnover and cytokine signaling.

**Conclusions:** Ganglioglioma represents a hybrid glioneuronal neoplasm in which developmental neuronal programs are co-opted by tumor-associated stromal and immune cues. This convergence establishes a permissive oncogenic niche that sustains precursor-like tumor cells and provides a mechanistic basis for both the tumor’s benign growth and its intrinsic epileptogenicity.

**Key Points:**

- Spatial transcriptomics identifies reproducible transcriptional programs that define the ganglioglioma microenvironment.
- Tumor-associated regions show transcriptional programs consistent with immature neuronal states together with ECM remodelling and immune activity.
- Single-cell reference data indicate that immature neuronal programs in ganglioglioma resemble radial glia-derived developmental states.

**Importance of the Study:** Ganglioglioma is a low-grade glioneuronal tumor that combines benign growth with pronounced epileptogenicity, yet the molecular basis of this dual behavior remains poorly understood. Through spatial transcriptomics integrated with single-cell analysis, we reveal that ganglioglioma architecture is defined by two interacting transcriptional axes: a residual glioneuronal network and a tumoral niche enriched for extracellular-matrix, vascular, and immune programs. Within these niches, immature neuronal-like tumor cells persist in a developmentally arrested state maintained by ECM-immune signaling. This spatially organized interplay between physiological and pathological programs explains both the low oncologic aggressiveness and high excitability of these lesions. Our findings provide molecular signatures that may refine diagnostic classification within the LEAT spectrum, delineate epileptogenic zones, and identify candidate pathways for therapeutic modulation of the ganglioglioma microenvironment.

## Introduction

Gangliogliomas (GGs) are rare, low-grade glioneuronal tumors and a leading cause of focal drug-resistant epilepsy in children and young adults^1^. They account for 1–2 % of all central nervous system (CNS) tumors and typically arise in the temporal lobe, where early surgical resection is key to seizure control^2^. Although the oncologic burden of most GGs is low^3^ and surgical outcomes are favorable, a clinically relevant subgroup exhibits incomplete seizure control or tumor recurrence^4^. These lesions frequently display atypical, infiltrative growth patterns extending beyond the cortical surface and disregarding anatomical boundaries^4^. Diagnosis itself remains challenging because GGs belong to the spectrum of low-grade epilepsy-associated neuroepithelial tumors (LEATs), which share overlapping morphological and molecular features^5-7^. Recently described entities such as PLNTY and pediatric low grade glioma^8,9^, which typically harbors a BRAFV600E mutation and shows strong CD34 expression with a GG-like methylation profile, further complicate classification^10-12^. Beyond classification, the mechanisms underlying the epileptogenic potential of GGs remain poorly understood. A central debate is whether the neuronal component represents a neoplastic, tumor-derived lineage or dysplastic cortical neurons secondarily overgrown by neoplastic glial cells^7,11^. These diagnostic and biological ambiguities highlight the need to define ganglioglioma-specific transcriptional programs to refine molecular classification, identify biomarkers, and clarify how tumor architecture relates to epileptogenicity. Previous studies have provided important insights into the cellular composition of GGs^13,14^, but systematic efforts to resolve their spatial organization are missing. Spatial transcriptomics now enables transcriptome-wide mapping directly on histological sections, offering unprecedented insight into tumor-stroma interactions and microanatomical architecture^15-17^. Here, we applied array-based unsupervised spatially resolved transcriptomics to a cohort of epilepsy associated GGs to delineate their transcriptional and microenvironmental organization. Our analysis uncovers spatially organized niches composed of immature, radial-glia–derived neuronal-like tumor cells embedded in reactive stroma enriched for extracellular-matrix remodeling, immune infiltration, and vascular signaling. These findings define ganglioglioma as a hybrid glioneuronal neoplasm in which developmental and microenvironmental programs coexist to drive both benign growth and intrinsic epileptogenicity.

## Materials and Methods

### Ethics

Detailed information of the permissions is provided in the supplementary data.

### Data Set Collection

This work uses Visium data of a total of eight ganglioglioma samples of which five were obtained from other works and one was created from our group, see Supplementary Table 1. Visium data sets prepared from our group included GG1, GG3, GG4, GG5, GG6. Tumor samples were formalin-fixed, paraffin-embedded, and sectioned for hematoxylin and eosin (H&E) staining. Spatially resolved transcriptomics was performed using the 10x Genomics Visium platform according to the manufacturer’s protocol. For our Spatial Transcriptomics Experiments, the Visium Spatial Gene Expression Slides (PN-20000233) with 6.5×6.5mm capture areas, surrounded by the 8×8 mm fiducial frame, were used (see CG000408 | Rev A). Briefly, tissue sections were hybridized with spatially barcoded oligonucleotides and then subjected to reverse transcription, cDNA synthesis, and library preparation. Sequencing was performed on an Illumina sequencer, and raw reads were processed using the 10x Genomics Space Ranger software suite. Additionally, this work makes use of three ganglioglioma Visium data sets from published works, which comprise GG_HDB1 and GG_HDB2^17^ and GG_Ext3^13^. Furthermore, eight published Visium samples of non-tumorous human cortex were used and referred to as healthy cortex^18^. Each data set has been downloaded with the respective accession code as provided in the original publication.

### Data Preprocessing and Module Identification

Spatial expression data was processed with SPATA2 (v3.1.4)^15^ and Seurat (v5)^19^. Protein-coding genes shared across all samples were retained using biomaRt. Each dataset was normalized independently with Seurat::SCTransform, and integration anchors were identified from 1,500 highly variable features using Seurat::SelectIntegrationFeatures, PrepSCTIntegration, and FindIntegrationAnchors (nn.method = “rann”, k.anchor = 5). Datasets were integrated with Seurat::IntegrateData (k.weight = 50, dims = 1:30). Shared nearest neighbor clustering (Seurat::FindClusters) was performed across resolutions 0.6–1.5, and cluster stability was assessed using the adjusted Rand index. Cluster-level mean expression values were aggregated into a gene-by-cluster matrix for downstream analysis. To evaluate the success of horizontal integration in removing technical or patient-specific factors, the aggregated gene expression profiles of all sample-wise clusters were correlated and hierarchically clustered to create meta-clusters. The distribution of sample origin, BRAF V600E mutation status, and patient sex was tested across these meta-clusters. For each categorical variable, contingency tables were constructed, and uniformity was assessed using Fisher’s exact or chi-squared tests, depending on cell counts. Effect sizes were quantified using Cramer’s V to estimate the strength of association. Weighted gene co-expression network analysis^20^ (WGCNA) was applied to the gene-by-cluster matrix to identify modules of co-regulated genes. A soft-thresholding power ensuring approximate scale-free topology was used to construct a signed adjacency and topological overlap matrix. Modules were defined by dynamic tree cutting and merged based on eigengene similarity. For each module, an eigengene (first principal component) summarized module activity, and hub genes were determined by intramodular connectivity (top 100 by kWithin).

### Single-cell deconvolution with cell2location

Cell2location was used to infer the spatial distribution of cell types in ganglioglioma tissue sections^21^. A single-cell RNA-seq dataset from five annotated ganglioglioma samples served as reference^13^. Genes shared with Visium data were retained, highly variable genes were selected, and a regression model was trained for 250 epochs to estimate cell-type expression signatures. For each Visium sample, data were filtered, normalized, and modeled with Cell2locationModel. Convergence was assessed by ELBO loss, and posterior cell-type abundances were aggregated by cluster and correlated with WGCNA module eigengenes.

### Module annotation with Gene Ontology (GO)

Hub genes were identified by quantifying each gene’s intramodular connectivity within its co-expression module. For each module, a topological overlap matrix (TOM) was computed from the weighted adjacency network using the same soft-thresholding power applied during WGCNA. The intramodular connectivity (k_(_Within_)_) of each gene was defined as the sum of its TOM connection strengths to all other genes in the same module. Genes were ranked by k_(_Within_)_, and the top 50 with the highest connectivity were designated as hub genes. Gene ontology enrichment was performed to annotate modules with functional interpretations using the the enrichGO() function from the R package clusterProfiler^22^ with ont=“BP” (Biological Processes) and pAdjustMethod=“BH” (Benjamini-Hochberg). Gene vectors corresponded to the hub gene lists of each WGCNA-module. Ontology terms with an adjusted p.value of lower than 0.05 were considered statistically significant.

### Module scoring

The activity of each WGCNA module was quantified at single-spot resolution by computing the difference between the mean expression of module genes and that of expression-matched control genes. For each module (Mk), the average log-normalized expression of all genes (g ∈ Mk) was contrasted against a control set (Ck) drawn from the same global expression distribution. All genes detected across ganglioglioma and healthy cortex samples were ranked by mean expression and divided into 30 quantile-based bins. For each gene g in Mk, up to 100 control genes were randomly sampled from the same bin, excluding all module genes. The module score was defined as the difference between the mean expression of Mk and Ck.

### Linear Mixed-Effects Modelling

Linear mixed-effects models were fitted using the R library lme4. Estimated marginal means and pairwise contrasts were obtained with emmeans, and *p*-values were adjusted using FDR correction. Module activity and spatial organization were compared between ganglioglioma and healthy cortex using models of the form *Score ∼ Tissue + (1* | *Donor)*, with donor as a random intercept to account for intra-donor correlation. For spatial organization, Moran’s I values (k = 6 nearest neighbors) quantified local clustering of module activity. To assess histological variation, models of the form *Score ∼ 0 + Histology + (1* | *Sample)* were fitted across annotated regions (tumoral, peritumoral, gray matter, white matter). Analyses were conducted on raw and z-scaled module scores.

### Processing and Analysis of Single Cells

Single-cell RNA-seq data (“Tumor 3” and “Tumor 5”) were processed with Seurat v5 using SCTransform for normalization and variance stabilization. Log_1+_-corrected residuals served as input for PCA, and the first ten components were used to construct shared nearest-neighbor and UMAP embeddings. Module activity was quantified per cell based on WGCNA programs, and high-scoring cells were identified using an elbow-point heuristic. Cell-type enrichment within modules was tested using Fisher’s exact test on 2×2 contingency tables, with *p*-values adjusted by the Benjamini–Hochberg FDR. Effect sizes were expressed as log_2_ enrichment ratios (*p*_(_observed_)_ / *p*_(_expected_)_), where positive values indicate enrichment and negative values depletion. Cells enriched for the black, pink, and turquoise modules were re-integrated using SCT-based integration with variance-selected features (excluding mitochondrial and ribosomal genes). Anchors were identified with FindIntegrationAnchors and corrected with IntegrateData^23^. The integrated data were clustered and visualized via PCA and UMAP. Marker genes were identified with the default settings of FindAllMarkers.

## Results

### Cohort characteristics and data integration

We analyzed a cohort of eight histologically confirmed GGs **(Fig. 1a)**. Five cases were collected at the Medical Center Freiburg, and three were obtained from previously published datasets^13,17^, of which two have not been analyzed with respect to ganglioglioma biology. All patients presented with drug-resistant epilepsy. Diagnoses were strictly established according to WHO CNS 2021 and ELAE criteria^3^: Each tumor exhibited a distinct ganglion-cell component, low proliferative index (Ki-67 < 5 %), and CD34 positivity by immunohistochemistry^24^. Targeted sequencing revealed a *BRAF* V600E mutation in five of eight cases, while three were wild-type (WT). All samples were processed using the 10x Genomics Visium platform **(Fig. 1a; Supplementary Fig. 1a)**.

**Figure 1.**
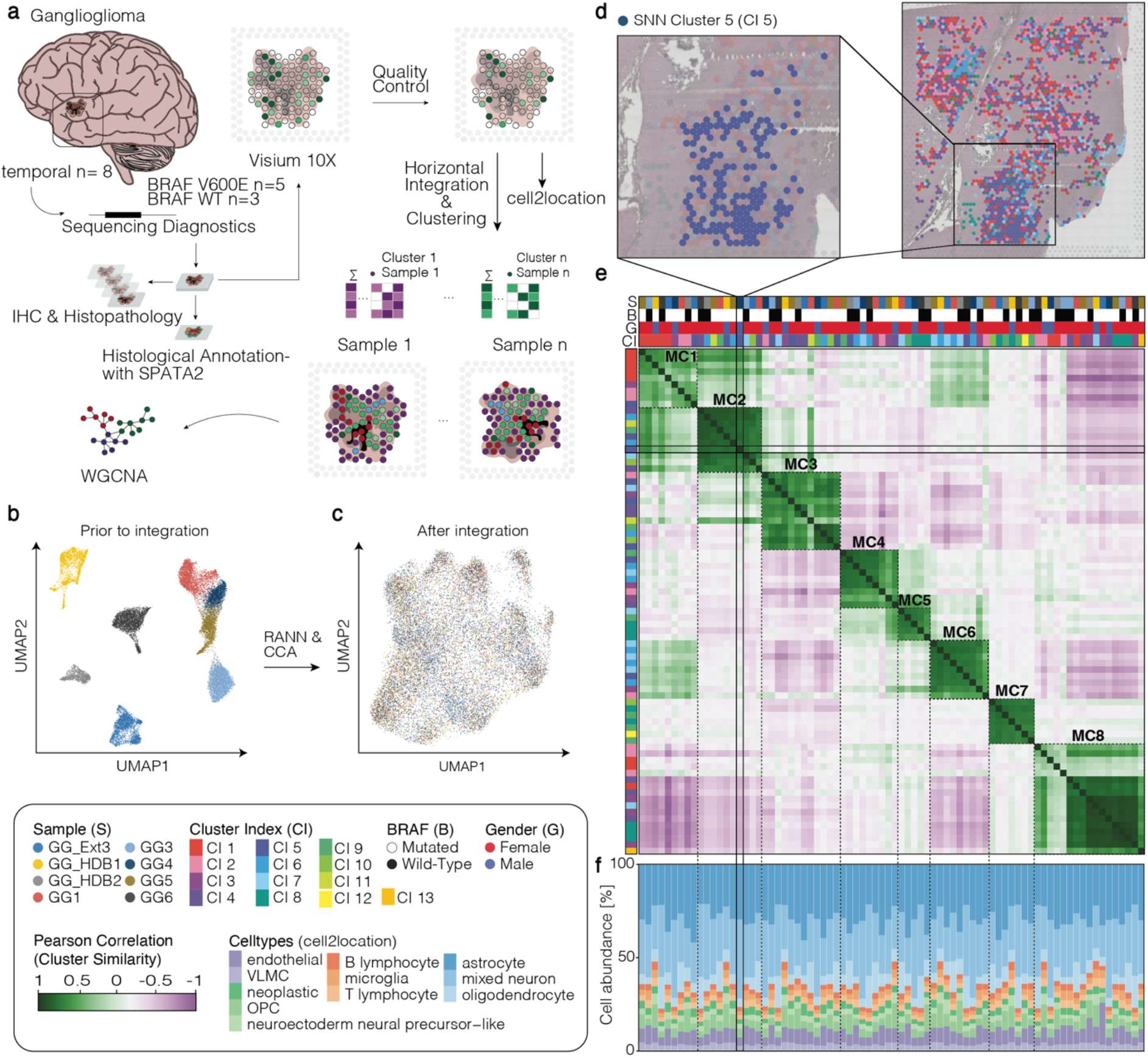
Data integration. a) Schematic overview of the study workflow. Eight histologically confirmed gangliogliomas (GGs; BRAF V600E, n = 5; wild type, n = 3) were analyzed by spatial transcriptomics (10x Visium). Following sequencing diagnostics, histopathologic validation, and annotation with SPATA2, datasets were integrated and analyzed using weighted gene correlation network analysis (WGCNA). b–c) Two-dimensional UMAP representation of all samples before (b) and after (c) horizontal integration using canonical correlation analysis (CCA) and approximate nearest-neighbor mapping, demonstrating alignment of shared transcriptional features across patients. d) Example of shared-nearest-neighbor (SNN) clustering in sample GG4. SNN cluster 5 (CI 5) is highlighted and visualized in spatial context, illustrating coherent regional grouping of transcriptionally similar spots. e) Correlation heatmap of sample-wise clusters, revealing eight reproducible meta-clusters (MC1–MC8) that define recurrent transcriptional domains within the ganglioglioma microenvironment. Metadata tracks indicate sample (S), BRAF status (B), and patient sex (G). f) Estimated cell-type composition of clusters based on reference deconvolution (Regal et al. 2023), showing consistent contributions of neuronal, glial, and immune populations across meta-clusters.

A primary objective of this work was to identify biological processes that characterize the microenvironment of epilepsy associated ganglioglioma and are consequently shared across a substantial number of samples. To mitigate tumor-intrinsic heterogeneity and batch-driven patient clustering **(Fig. 1b)**, we applied horizontal integration to align shared biological signals while minimizing technical noise. To this end, integration was performed with radius-based approximate nearest-neighbor mapping and canonical correlation analysis^23^ **(Fig. 1c)**. Samples were then re-split for sample-by-sample transcriptional clustering via Shared-Nearest-Neighbor. To ensure that our integration process succeeded, we assessed similarity of clusters across samples by correlating their gene-expression profiles, which was again hierarchically sorted to group clusters that are similar to each other in what they transcriptionally represent. **Figure 1e** displays the resulting correlation structure suggesting that there are eight meta-clusters in our ganglioglioma cohort, eight recurrent transcriptional domains. Meta data annotations for sample origin (S), *BRAFV600E* mutation status (B), and patient gender (G) are displayed at the top level of the heatmap and show comparable representation across meta-clusters. (Fisher’s exact test; all p > 0.99; Cramer’s V < 0.13.) This supported the effectiveness of our integration in minimizing patient-specific effects and enabling a unified analysis of ganglioglioma biology.

### Weighted gene co-expression analysis reveals eight recurrent transcriptional programs defining the ganglioglioma microenvironment

We next aimed to uncover the gene-level architecture underlying these patterns. Rather than continuing with empirical meta-clusters, we applied weighted gene co-expression network analysis (WGCNA) to the aggregated cluster-by-gene matrix to identify modules of coordinately expressed genes^20^. This network-based approach identified eight distinct transcriptional modules representing conserved biological programs across the ganglioglioma cohort. Each module was assigned a color for identification (**Fig. 2a)**. Module eigengenes were correlated with cell2location-derived cell-type abundance estimates to infer the dominant cellular contributors^21^ (**Fig. 2b)**. Hub genes were identified, and functional annotation via gene ontology (GO) enrichment^22^ provided biological interpretation **(Supplementary Figs. 2–4)**. Collectively, modules could be categorized into three functional groups: (i) preserved cortical organization (blue, brown, and red), (ii) glial responses (green and yellow), and (iii) tumor associated programs (turquoise, black, and pink). The blue module (N_Genes_=177) contained synaptic genes such as *SNAP25* and *GRIN1*, and mapped to GO terms related to synaptic signaling (GO:0099504, FDR < 0.01). The brown module (N_Genes_=168) was associated with excitatory neuronal markers (*CAMK2N1, CAMK2A, GAP43*) and was linked to axonal transport (GO:0098930, FDR < 0.01). Together, blue and brown primarily reflected mature neuronal biology (**Fig. 2e)**. The red module (N_Genes_=129) was dominated by oligodendrocyte markers like *PLP1* and *MBP* **(Fig. 2f**). GO analysis was consistent with active myelination and oxygen dependent energy metabolism (GO:0032291, GO:0015980, all FDR < 0.05). The yellow module (N_Genes_=152) represented a reactive, progenitor-like state enriched for *PDGFRA, GFAP* and *ALDOC* (**Fig 2c)** and was functionally linked to precursor proliferation and gliosis (GO:0042063, GO:2000179, all FDR < 0.05). The green module (N_Genes_=138) reflected a homeostatic glial program characterized by genes like *SLC1A3* and *KCNJ10* (**Fig. 2d**) and was linked to neurotransmitter uptake and metabolic support (GO:0098712, GO:0010975, all FDR < 0.01). The turquoise module (N_Genes_=361) was enriched for genes involved in vascular remodeling and extracellular-matrix reorganization like *VEGFA, ACTA2, LTBP3, COL3A1, FN1* alongside immune mediators *CD74, B2M, CXCL12* (**Fig. 2h)**. The black module (N_Genes_=58) comprised stress-response and cell-matrix-interaction genes like *CD44, CD63, CHI3L1* **(Fig.2i)**. GO analysis mapped it to adhesion (GO:0007160, FDR < 0.05) and neuron projection regeneration (GO:0031102, FDR < 0.01). The pink module (N_Genes_=55) contained developmental and neuronal-signaling genes like *HOPX, CHL1, NRCAM, GRIA1, GRIA3, APP* **(Fig. 2g)**. GO analysis highlighted enrichment for synapse organization (GO:0050808, FDR < 0.01), axonogenesis (GO:0007409, FDR < 0.01) next to stem-cell differentiation (GO:0048863, FDR < 0.01).

**Figure 2.**
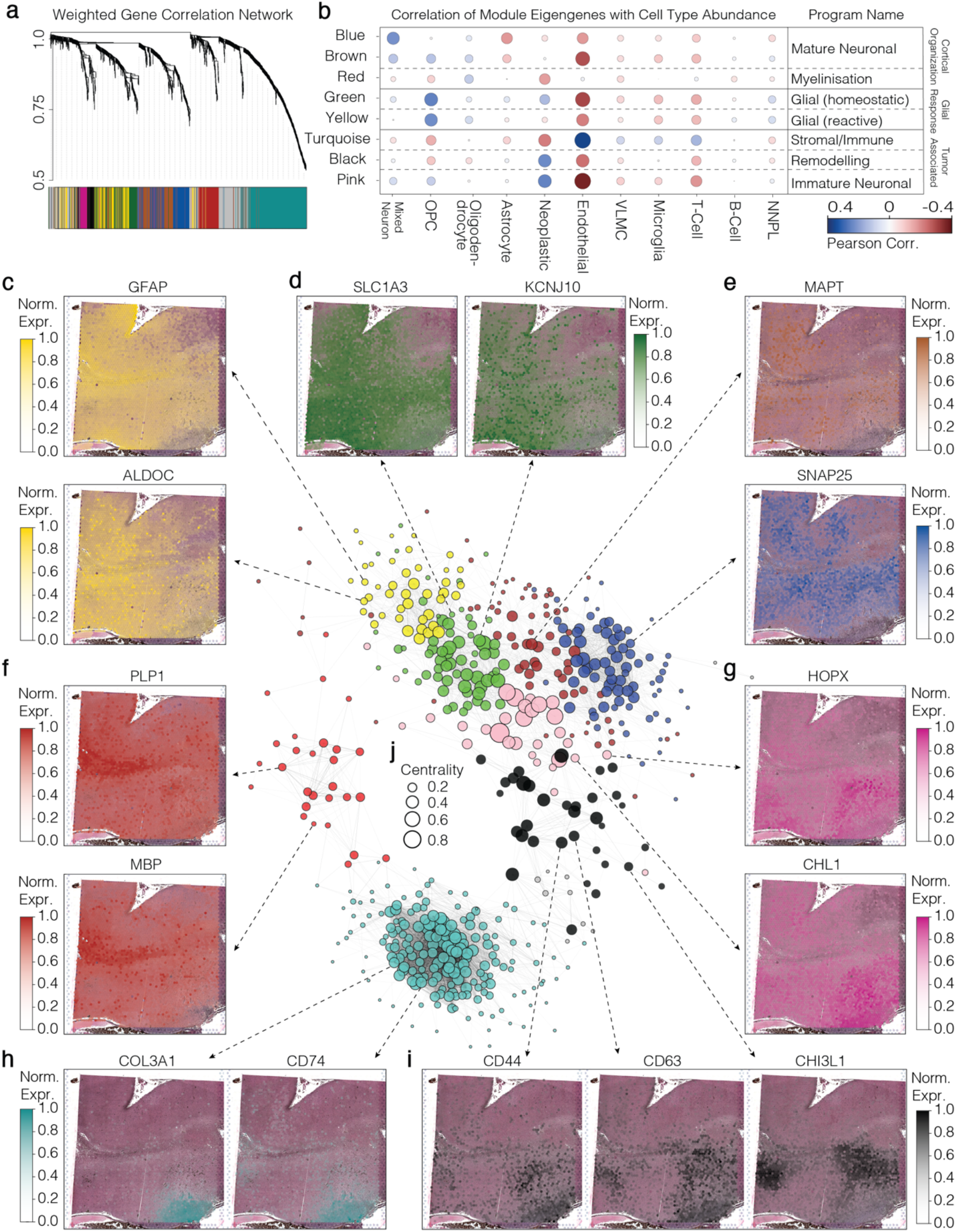
The transcriptional landscape of the ganglioglioma microenvironment revealed by WGCNA. **a)** Hierarchical clustering dendrogram of gene modules identified. **b)** Correlation matrix showing the association between module eigengenes and cell-type abundance scores derived from cell2location using the annotations of Regal et al. 2023. **c–i)** Spatial expression levels of representative genes on sample GG5 exemplifying characteristic biological processes of each module.: **c)** reactive astrogliosis, **d)** homeostatic glial support, **e)** mature neuronal activity, **f)** oligodendrocytic myelination, **g)** progenitor-like/neurodevelopmental **h)** immunity as well as stromal and vascular remodeling, **i)** ECM remodeling and tumor-intrinsic immune-like signaling. **j)** Subnetwork of genes used for WGCNA visualized as a correlation network and colored by gene-module assignment.

### Disentangling physiological from pathological processes in ganglioglioma

So far, we identified recurrent transcriptional programs in our ganglioglioma cohort, several of which likely reflect pathological processes. However, gene ontology analysis indicated that the blue, brown, and red modules captured processes characteristic of the healthy neocortex, suggesting that a subset of Visium spots in the GG cohort represented intact or only mildly infiltrated cortical tissue. To resolve whether the identified transcriptional programs reflect ganglioglioma-specific or physiological cortical biology, we established a healthy reference dataset consisting of eight dorsolateral prefrontal cortex samples^18^, derived from non-neoplastic human brain tissue. Program activity for each spot in both cohorts was quantified using an expression-matched scoring approach that compared module gene expression with randomized background sets. A score of zero indicated no deviation from background, while positive values reflected increasing activation strength **(Fig 3d-g, Supplementary Fig. 6a-d)**. This enabled direct comparison of program activity between healthy and ganglioglioma tissues. Statistical effects were estimated using likelihood-ratio tests from linear mixed models with donor as a random intercept **(Supplementary Fig. 5a–b)**. Programs associated with mature neuronal function showed significantly higher activity in healthy cortex (β = -0.11 to -0.22, all FDR < 0.05). Myelination did not show a significant difference between cohorts (β = 0.03, FDR = 0.56). In contrast, glial response and tumor associated programs were significantly upregulated in ganglioglioma tissue (β = 0.12 to 0.24, all FDR < 0.01). Next, we aimed to quantify where program activity above baseline reflected genuine biological organization. Because Visium spots capture transcripts from multiple neighboring cells, the resulting modules reflect coordinated transcriptional programs within local tissue regions rather than discrete cellular populations. Therefore, we reasoned that genuine activation should manifest as spatially coordinated upregulation. We computed spatial autocorrelation (Moran’s I) for each program and compared values between cohorts **(Supplementary Fig. 5c–d)**. Programs related to cortical organization showed comparable spatial coherence across cohorts (β = – 0.06 to –0.02, all FDR > 0.2). In contrast, glial response and tumor associated programs exhibited markedly stronger spatial organization in ganglioglioma (β = 0.20–0.30, all FDR < 0.005). Together, these analyses demonstrate that while elements of normal cortical biology persist within ganglioglioma, reactive glial and tumor associated processes display both higher activity and enhanced spatial organization, defining the pathological transcriptional architecture of the lesion.

**Figure 3.**
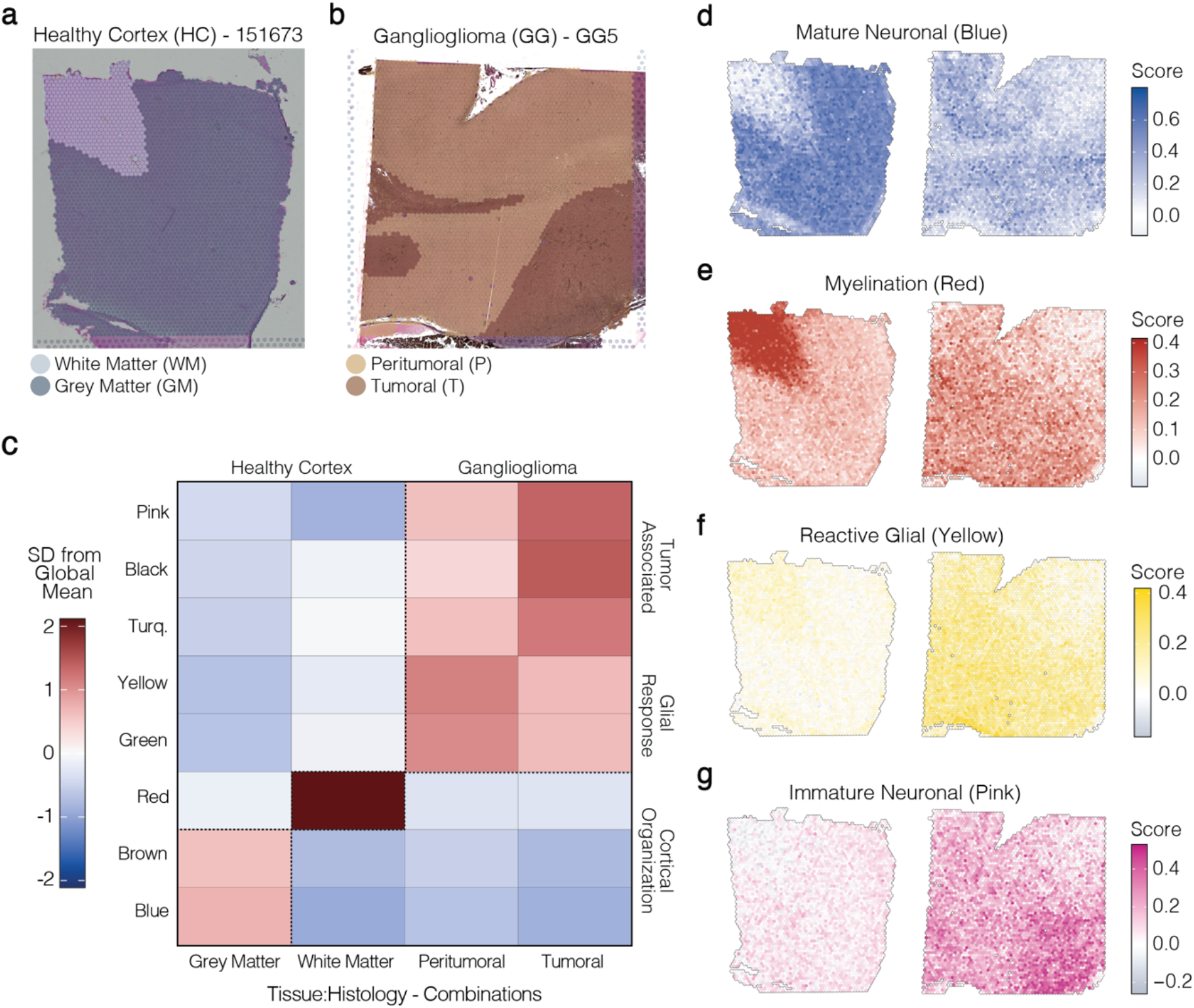
Disentangling physiological and pathological transcriptional programs in ganglioglioma. **a-b)** Representative examples of the control cohort of healthy cortices (a) and ganglioglioma (b) colored by the histological classification. **c)** Heatmap of standardized marginal means as estimated by our mixed effect linear modelling of module activity across tissue and histology. **d-g)** 2D Surface plots contrast module expression and spatial organization of microenvironmental niches in healthy cortex (left column) and ganglioglioma (right column). The color mapping is scaled to the global range of module activity in both cohorts highlighting how ganglioglioma feature lower (diluted) expression of healthy cortical modules (d-e) and higher expression of glial and tumorous modules (f-g).

### Spatially organized transcriptional domains reveal coordinated pathological programs in ganglioglioma

We next examined how program activity relates to histological architecture. Despite limited image resolution, two ganglioglioma samples displayed visually distinct regions corresponding to mildly infiltrated (peritumoral) and infiltrative or subpial (tumoral) growth **(Supplementary Fig. 7a–b)**. Program activity was modeled across gray and white matter in healthy cortex and across peritumoral and tumoral regions in ganglioglioma **(Fig. 3a–b)**. Healthy cortical function exhibited pronounced gray–white matter separation in healthy cortex (β = –2.35 to 1.73; all FDR < 0.01), confirming that the respective programs reflect healthy cortical organization. Glial response was elevated in ganglioglioma, most prominently in peritumoral regions, whereas tumor associated programs showed a progressive increase from peritumoral to tumoral tissue (β = –1.09 to –0.63, all FDR < 0.01). To further explore inter-program relationships, we correlated spot-level program scores across all ganglioglioma samples. The resulting correlation structure **(Fig. 4a)** combined with the region-resolved modeling **(Fig. 4b)** suggested that the ganglioglioma microenvironment is organized into two functionally distinct transcriptional domains. The first defines a glioneuronal domain **(Fig. 4c)** composed of mature neuronal processes that are co-expressed with glial response. The second defines a tumoral domain formed by the pink, black, and turquoise modules, characterized by high inter-module correlation and strong upregulation within histologically disrupted regions **(Fig. 4b, d)**. Their coordinated activation links immature neuronal, matrix-remodeling, and vascular-immune processes into a coupled transcriptional network. Together, these results indicate that ganglioglioma is a spatially hybrid tissue in which a residual glioneuronal framework provides the structural substrate for embedded tumoral niches.

**Figure 4.**
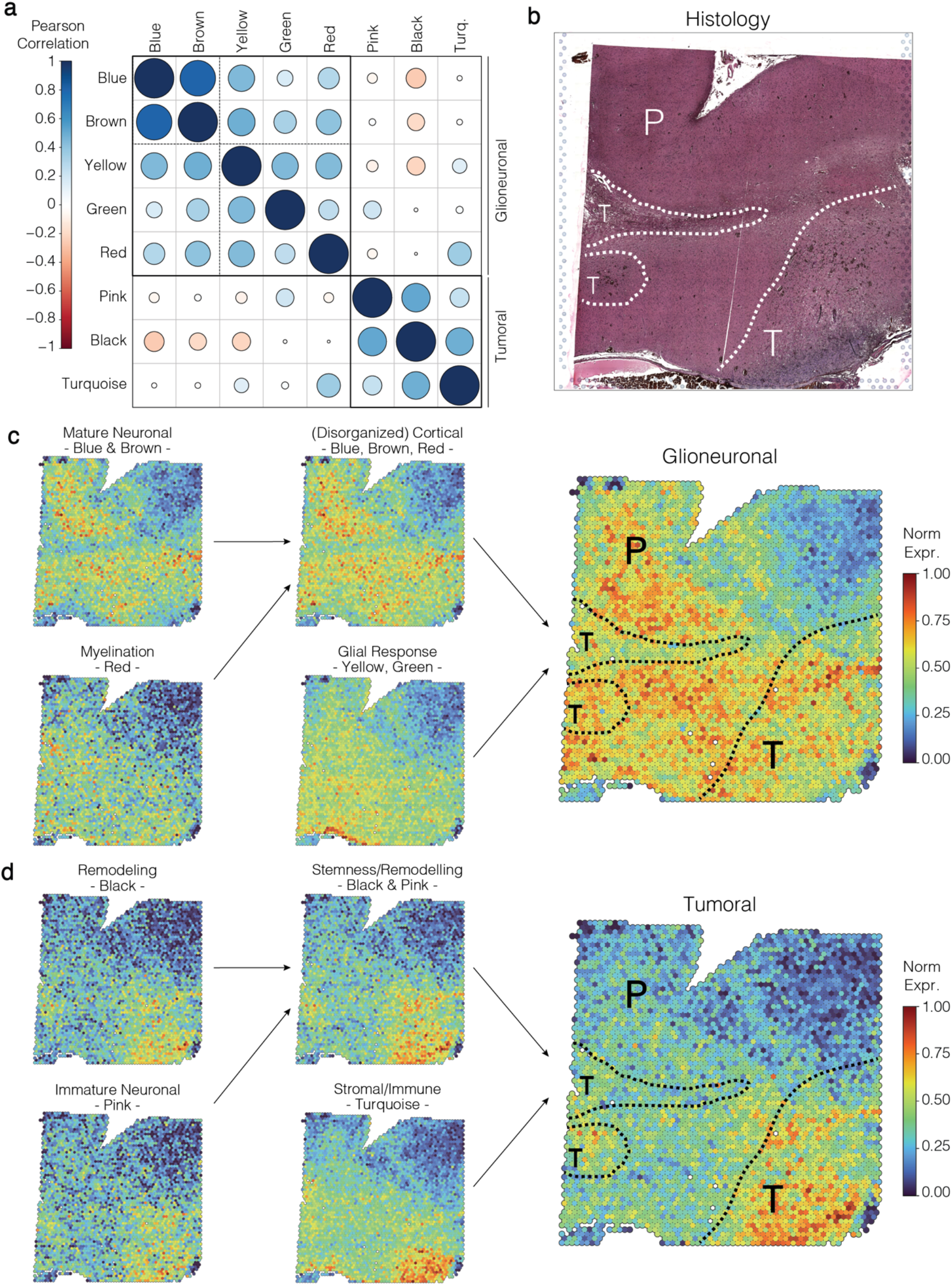
The transcriptional architecture of the ganglioglioma microenvironment. **a)** Spot-level correlations among all eight WGCNA modules across the ganglioglioma cohort reveal two principal transcriptional axes. **b)** Dotted lines indicate the histological boundary between peritumoral and tumoral compartments in sample GG5. **c–d)** Hierarchical representation of how microenvironmental niches within ganglioglioma assemble into a higher order organization defined by two major transcriptional axes and their relationship to histological structure..

### Single-cell analysis defines an immature neoplastic neuronal core within reactive ganglioglioma niches

The identification of distinct functional domains raised the question of which cellular states and tissue components give rise to each transcriptional organization. To resolve this, we leveraged the pre-labeled single-cell RNA-sequencing dataset13 (Fig. 5a) and scored every single cell for the eight modules identified in our spatial analysis. For each module, an elbow heuristic defined the top fraction of cells contributing most strongly to its biology (Supplementary Fig. 8a). The resulting cellular composition closely mirrored the functional annotation derived from the spatial data. The mature neuronal programs were dominated by mixed neuronal populations (log_2_ enrichment = 2.7–3.1; all FDR < 10^−18^). The myelination program was enriched in oligodendrocytes (log_2_ enrichment = 1.6; FDR = 1.7 × 10^−159^). The glial response program showed peak activity in OPC- and neuroectoderm-like precursor cells (log_2_ enrichment = 2.1–2.5; all FDR < 10^−20^). The stromal and immune program was primarily composed of microglia and endothelial cells (log_2_ enrichment = 1.8–2.3; all FDR < 10^−15^). The immature neuronal and remodelling programs consisted almost exclusively of neoplastic cells (log_2_ enrichment = 1.1–1.4; all FDR < 10^−32^). We next focused on the cellular architecture underlying the tumoral domain. Cells scoring highest for these programs were pooled, subset, and re-integrated for higher-resolution analysis (Fig. 5f). Single-cell clustering yielded six distinct clusters (C2–C7; C1 was excluded due to incoherent transcriptional identity, see Methods). The mapping from original subtype annotations to new clusters is shown in Fig. 5g. Marker-gene detection for cluster characterization was performed using Wilcoxon rank-sum testing (Supplementary Fig. 8c). A comprehensive list of significant marker genes for all clusters is provided in the Supplementary Data. This analysis revealed that clusters C2, C4, and C6 represented transcriptionally distinct yet related neoplastic populations. C2 showed a coherent signature co-expressing immature excitatory neuronal genes HOPX (log_2_FC = 0.9, FDR < 0.001), MAP2 (log_2_FC = 0.7, FDR < 0.001), and NLGN1 (log_2_FC = 0.5, FDR < 0.001) together with radial-glia regulators CDH2 (log_2_FC = 0.5, FDR < 0.01) and TNR (log_2_FC = 0.7, FDR < 0.001), defining a hybrid neuronal–glial progenitor state. C4 shared this immature background but showed stronger MAP2 expression (log_2_FC = 1.1, FDR < 0.001), while C6 retained NLGN1 expression (log_2_FC = 0.4, FDR < 0.05) alongside extracellular-matrix remodeling genes. The remaining clusters reflected the reactive microenvironment: C3 corresponded to activated microglia (CX3CR1, log_2_FC = 8.8, FDR < 0.001), C5 to oligodendrocytic cells (SOX2, log_2_FC = 2.3, FDR < 0.001), and C7 to endothelial cells (CDH5, log_2_FC = 6.7, FDR < 0.001). In summary, clusters C2, C4, and C6 likely represent a continuum of neuronal differentiation, ranging from radial glia-like progenitors to immature neurons. These results indicate that ganglioglioma is organized around an immature neuronal core embedded within reactive glial, immune, and vascular compartments, providing the cellular basis for the hybrid architecture observed in spatial data.

**Figure 5.**
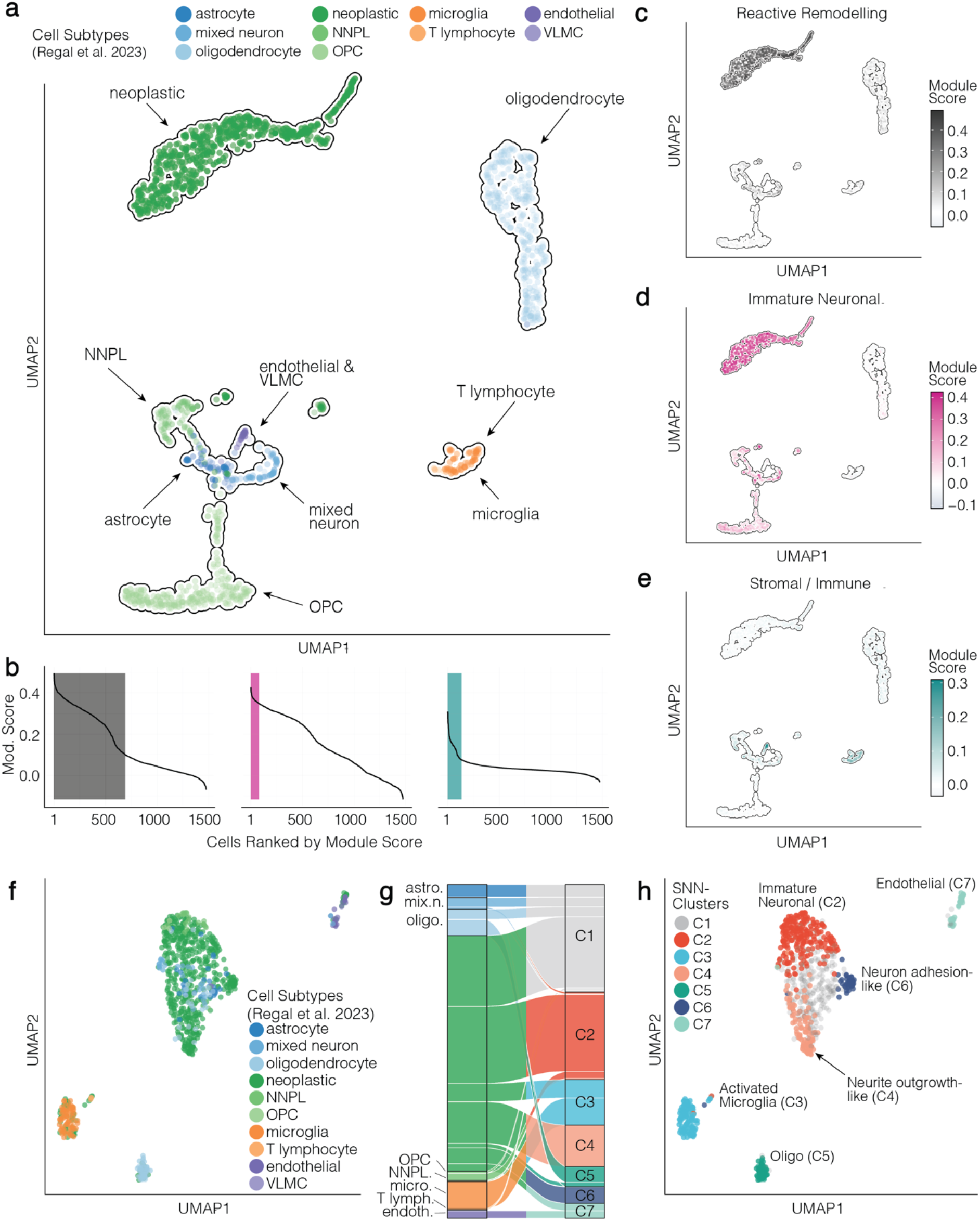
Single-cell composition of ganglioglioma neoplastic microenvironments. **a)** UMAP visualization of single cells from Tumors 3 and 5. b) Cells ranked by neoplastic module score; shaded areas denote thresholds for downstream inclusion as determined by elbow identification. **c–e)** UMAPs as in (a), colored by module scores, illustrating strong overlap of high spatial transcriptomic-derived neoplastic module scores with neoplastic cell types (c, d) and additional enrichment in endothelial and immune-associated niches (e). **f)** UMAP of the selected subset of cells following horizontal integration across samples. **g)** Sankey-plot depicting label transfer from initial cell subtype annotation to newly derived SNN clusters. **h)** UMAP from (f) showing the final cluster identities used for downstream analyses, with neoplastic cells distributed across three lineage-related programs (C2, C4, C6) and additional endothelial, oligodendrocytic and microglial compartments.

## Discussion

In this study, we applied spatial transcriptomics and mapped the molecular organization of ganglioglioma. We identified a spatial architecture defined by two distinct transcriptional domains. The glioneuronal domain was characterized by preserved yet disorganized programs of mature neuronal activity and myelination, accompanied by reactive gliosis, consistent with previous reports^13^. These findings indicate that elements of physiological cortical organization persist within ganglioglioma in a disrupted and reactive state. In contrast, the tumor associated domain comprised transcriptional programs related to immature neuronal signaling, extracellular matrix remodeling, and vascular–immune activity. These were enriched in histologically disturbed tissue regions. Among these, the pink module was enriched for neuron-projection guidance and glutamate receptor signaling alongside stemness-associated pathways and likely reflects an immature neuronal transcriptional state. This program contrasts with the mature neuronal activity observed in the glioneuronal domain and supports the presence of immature neuronal-like networks within ganglioglioma. This framework is consistent with gangliogliomas frequent association with epilepsy.

At the single-cell level, the transcriptional continuum identified within the tumor associated domain also points to immature neuronal-like states and resembles early developmental programs rather than purely proliferative tumor populations. These states showed similarity to radial glia due to the combination of early progenitor signatures (HOPX, TNR, CDH family), HOPX, Tenascin-C and neuronal markers (MAP2, GRIA1, GRIK1). This suggests the presence of a transcriptional cellular state that combines precursor-associated and neuronal features without full maturation. These findings are broadly consistent with prior work, particularly Regal et al., who also described immature neuronal or neuroectodermal stem cell– like populations in ganglioglioma. Differences in cohort composition and diagnostic criteria, however, may influence the interpretation of these states. While Regal et al. included a less strictly selected cohort with regard to proliferative activity, our study focuses on a clinically and histopathologically strictly defined cohort of epilepsy-associated gangliogliomas fulfilling WHO 2021 criteria. We interpret our findings within a developmental framework, suggesting persistence of early neuronal precursor-like programs rather than reversion to an immature state. This interpretation is supported by the characteristic mismatch between early childhood onset and slow tumor growth kinetics. With proliferation indices typically below 5%^3^ and often near 1%^25^, these tumors expand slowly. Their frequent presentation in childhood suggests that part of the tumor mass may arise during earlier stages of brain development when radial glia are active. Notably, our data do not permit lineage inference at single-cell resolution, and further studies will be required to resolve this question.

To our knowledge, our cohort represents the largest to date in which spatial transcriptomics has been used to investigate the microenvironment of ganglioglioma. Nevertheless, the sample size remains limited. To mitigate these limitations, we implemented a conservative analytical strategy. Horizontal integration was applied across samples from three institutions to align shared microenvironmental modules despite the risk of signal collapse^13,25,26^. Moreover, functional annotation was confined to hub genes to reduce statistical noise and focus on main transcriptional drivers. Future studies using higher-resolution, multimodal, or lineage-tracing approaches will be required to further define the cellular origin and the mechanisms underlying our observations. Moreover, it would be valuable to investigate whether the immature neuronal program we assume to be involved in the formation of pathological neuronal circuits is specific to gangliogliomas that present with seizures. Comparing its activity between tumors with and without epileptogenicity could provide further insight into its role in the manifestation of this disease.

In summary, our findings support a model of ganglioglioma as a developmental hybrid tumor organized around immature, radial-glia-derived neuronal-like cells embedded in a reactive cortical microenvironment **(Supplementary Fig. 9)**. This hybrid architecture may underlie both the benign oncological course and the pronounced epileptogenic potential that define a tumor entity that remains incompletely understood.

## Ethics

The study design, data evaluation, and imaging procedures were given clearance by the ethics committee at the University of Freiburg, as delineated in protocols 100020/09 and 472/15_160880. All methodologies were executed in compliance with the guidelines approved by the committee. Informed consent, in written form, was received from all participating subjects. The Department of Neurosurgery of the Medical Center at the University of Freiburg, Germany, was responsible for securing preoperative informed consent from all patients participating in the study.

## Authorship

The study was designed and coordinated by DHH and DD. Analysis was conducted by JK and LH. Main part of the manuscript was written by JK and LH. Method part of the manuscript was written by JK and JM. Main figures were created by JK. Supplementary figures were created by JK. GG samples were selected and diagnosed by RS. Visium samples derived from Freiburg were prepared by JM and DNZ. The manuscript was edited by OS, JB, JZ, JKB, PS and FS.

## Data Availability

Raw and processed data will be made available upon publication.

## Acknowledgements

ChatGPT (OpenAI) was used for language editing. All scientific content, analyses, and interpretations were conducted by the authors. The authors take full responsibility for this work.

